# Proprioceptive Deficits in Inactive Older Adults are not Reflected in Discrete Reaching Performance

**DOI:** 10.1101/393785

**Authors:** Nick M. Kitchen, R Chris Miall

**Affiliations:** School of Psychology, University of Birmingham, Birmingham, UK.; Dept. of Speech & Hearing Science, University of Washington, Seattle, WA, USA.

**Author notes:** **Address for correspondence:** Nick Kitchen, Dept. of Speech & Hearing Science, 1417 NE 42nd St, Seattle, WA, 98105.

**Keywords:** Proprioception, Ageing, Reaching, Sensorimotor Control

## Abstract

During normal healthy ageing there is a decline in the ability to control simple movements, characterised by increased reaction times, movement durations and variability. There is also growing evidence of age-related proprioceptive loss which may contribute to these impairments. However this relationship has not been studied in detail for the upper limb. We recruited 20 younger adults (YAs) and 31 older adults (OAs) who each performed 2 tasks on a 2D robotic manipulandum. The first assessed dynamic proprioceptive acuity using active, multi-joint movements towards visually presented targets, with movement constrained by the robot to a predefined path. Participants made perceptual judgements of the lateral position of the unseen arm. The second was a rapid motor task which required fast, accurate movements to the same targets in the absence of hand position visual feedback, and without constraint by the robot. We predicted that the variable proprioceptive error (uncertainty range) from Task 1 would be increased in physically inactive OAs and would predict increased movement variability in Task 2. Instead we found that physically inactive OAs had larger systematic proprioceptive errors (bias). Neither proprioceptive acuity nor bias was related to motor performance in either age group. We suggest that previously reported estimates of proprioceptive decline with ageing may be exaggerated by task demands and that the extent of these deficits is unrelated to discrete, ballistic movement control. The relationship of dynamic proprioceptive acuity with movement control in tasks which emphasise online proprioceptive feedback for performance is still unclear and warrants further investigation.

## Introduction

As we get older there is a general decline in motor system physiology which affects the ability to perform simple movements. This includes degradation of musculature through loss and remodelling of muscle motor units (Lexell, 1995; Morley, Baumgartner, Roubenoff, Mayer, & Nair, 2001; Slack, Hopkins, & Williams, 1979), as well as degeneration of efferent peripheral nerves and the neuromuscular junction (Ceballos, Cuadras, Verdu, & Navarro, 1999; Jacobs & Love, 1985; Valdez et al., 2010) which disrupts transmission of motor commands and impairs the ability to perform movements as intended. This is characterised in advanced age by increased movement duration (Contreras-Vidal, Teulings, & Stelmach, 1998; Helsen et al., 2016; Ketcham, Seidler, Van Gemmert, & Stelmach, 2002), as well as increased spatial (Darling, Cooke, & Brown, 1989; Seidler, Alberts, & Stelmach, 2002) and temporal (Contreras-Vidal et al., 1998; Yan, Thomas, Stelmach, & Thomas, 2000) variations during a range of different movement tasks. Interestingly, this is often coupled with a maintenance of endpoint accuracy (Helsen et al., 2016; Lee, Fradet, Ketcham, & Dounskaia, 2007; Seidler-Dobrin & Stelmach, 1998) which is thought to be achieved through increased movement duration, reaction time and by online corrective mechanisms which are frequently observed in this population (Helsen et al., 2016; Ketcham et al., 2002).

In addition to motor physiology, loss of proprioception has also been suggested as a contributing factor to the presentation of these age-related motor deficits. Specifically, there is growing evidence to show decline of this sensation through a range of different measurement techniques (see Goble, Coxon, Wenderoth, Van Impe, & Swinnen, 2009 for review), including limb position matching to both passively (Adamo, Alexander, & Brown, 2009; Adamo, Martin, & Brown, 2007; Helsen et al., 2016; Herter, Scott, & Dukelow, 2014; Lei & Wang, 2018) and actively (Schaap, Gonzales, Janssen, & Brown, 2015) derived reference positions. Age-dependent deficits have also been reported in thresholds for detecting passive joint displacement (Helsen et al., 2016; Wright, Adamo, & Brown, 2011) and in two alternative forced-choice paradigms involving position estimates of active, multi-joint movements (Cressman, Salomonczyk, & Henriques, 2010). This age-related loss of acuity appears to be amplified by physical inactivity (Adamo et al., 2009; Helsen et al., 2016; Wright et al., 2011) and in the lower limb, these deficits have been associated with impairments in functional motor measures including balance, posture, mobility and incidence of falls (Hurley, Rees, & Newham, 1998; Lord, Clark, & Webster, 1991; Sorock & Labiner, 1992; Wingert, Welder, & Foo, 2014). In spite of these reports, the extent to which proprioceptive loss contributes to age-related movement deficits of the upper limb is still poorly understood.

Recently, Helsen et al. (2016) attempted to address this by associating measures from two passive proprioceptive assessment techniques with participants’ performance in rapid, target-based wrist movements. Similar to previous reports, they found physically inactive older adults had prolonged detection thresholds for passive wrist displacement and increased matching errors to passively defined reference positions, indicating loss of proprioceptive acuity. But despite reporting stereotypical age-related motor kinematic impairments, the authors did not find an association between proprioception and motor performance. From this, they concluded that proprioceptive impairments can be overcome in ageing by greater reliance on predictive, feed-forward mechanisms of motor control.

However, since limb position sense can be directionally modulated by corollary discharge (Smith, Crawford, Proske, Taylor, & Gandevia, 2009), the proprioception experienced during active, voluntary movement is likely different to that of passive displacements. Indeed, active movement to participant-defined reference positions has been shown to reduce position matching errors compared to traditional, passive methods in both younger (Erickson & Karduna, 2012; Lönn, Crenshaw, Djupsjöbacka, Pedersen, & Johansson, 2000) and older (Langan, 2014) adults, demonstrating how sense of effort affects performance on these tasks. Hence, the null relationship of upper limb proprioception and motor control reported by Helsen et al. (2016) may actually reflect the difference in proprioceptive perception between passive and active movement. Furthermore, impairments in working memory and attention have been shown to confound position matching errors in ageing (Boisgontier, Olivier, Chenu, & Nougier, 2012; Goble, Mousigian, & Brown, 2012), which further advocates the use of alternative proprioceptive acuity assessments for investigating an association with voluntary movement control in this population.

Yet reports directly comparing age groups on active movement-based proprioceptive tasks which limit dependence on working memory are scarce. Cressman et al. (2010) measured shifts in sensed limb position associated with adaptation of reaches to a visual rotation in a group of older and younger adults. Sensed limb position was assessed by asking participants to make active, multi-joint reaching movements constrained to a tight, pre-defined trajectory, before making instantaneous judgements of their unseen limb relative to a visually presented reference position. These two-alternative forced choice responses were then gathered and used to estimate both systematic (bias) and variable (uncertainty range) proprioceptive errors; only the latter showed age-related increase, with marginal statistical significance. Variants of this task have been reported elsewhere (Cressman & Henriques, 2009; Ostry, Darainy, Mattar, Wong, & Gribble, 2010), but this was the first report of its use with an ageing population. Critically, since this type of task reduces dependence on working memory and utilizes active movements, it may be more suited for the investigation of age-related proprioceptive loss and voluntary movement control. Moreover, if it is indeed the case that proprioceptive uncertainty increases with ageing, then this elevated sensory noise could make the sensory consequences of motor commands unpredictable (Miall & Wolpert, 1996) and thus lead to more variable movement characteristics, which are frequently reported for the older adult population (Darling et al., 1989; Ketcham et al., 2002; Seidler et al., 2002). As such, the proprioceptive uncertainty estimate derived from this type of task makes for a compelling predictor of motor performance in the ageing population.

The aim of this experiment was therefore to assess, in groups of older and younger adults, the extent to which dynamic, multi-joint proprioceptive acuity of the upper limb could predict performance on a fast, targeted reaching movement task. We predicted that physically inactive older adults would exhibit larger proprioceptive uncertainty ranges and that this would predict greater variation in motor performance. Conversely, since a systematic perceptual error (assessed as proprioceptive bias), may be easier to predict and account for during motor control, we predicted bias would be unrelated to motor performance for either age group.

## Methods

### Participants

Thirty one older adults (OAs) aged 65 years or older (11 male, 71.2 ± 4.5 yrs), and 20 younger adults (YAs) aged 18-25 years (11 male, 20.4 ± 2.0 yrs) participated in the experiment after giving informed consent; the University of Birmingham ethics panel approved the study. All participants were right-hand dominant as defined by a laterality quotient of 30 or higher on the 10-item Edinburgh Handedness Inventory (Oldfield, 1971). Participants were excluded if they had any history of neurological illness, or carpal tunnel syndrome, arthritis or similar movement pains or limitations in the arm, wrist or fingers. OAs also completed the Montreal Cognitive Assessment (MoCA) and were only included in the analysis if they scored 26 or above out of 30, which is considered to indicate normal cognitive functioning (Nasreddine et al., 2005).

### Experimental Set-Up

Participants sat in front of a 2D-planar robotic manipulandum (vBOT; Howard, Ingram, & Wolpert, 2009) which provided a low-inertia, low-friction means of recording simple reaching movements in a 40×64cm workspace (Figure 1A). With their foreheads resting against a padded metal frame approximately 10cm behind the edge of the workspace, participants grasped the manipulandum handle with their right hand and were asked to look down onto a mirrored surface. This blocked direct view of the hand and arm and reflected images from a large, horizontally mounted monitor display. Target locations and visual feedback of hand position were presented in this way, with the cursor (when displayed) spatially coincident with the centre of the vBOT handle. Recordings of the vBOT handle position were sampled at 1kHz with any applied forces updated at the same rate. In both the dynamic proprioceptive and rapid motor reaching tasks, participants made reaching movements from a white 1cm radius start position located 8cm into the workspace (approximately 28cm from the participant’s torso). Participants made reaching movements to one of three positions, shown by a 1cm radius grey target, which were located 20cm from the start position at 30°, 90° and 150° elevation (Figure 1B). When made available, hand position feedback was provided on a real-time basis by a 0.5cm radius white cursor that was always spatially congruent with the vBOT handle. In all cases targets were presented in a pseudorandomised order.

**Figure 1:**
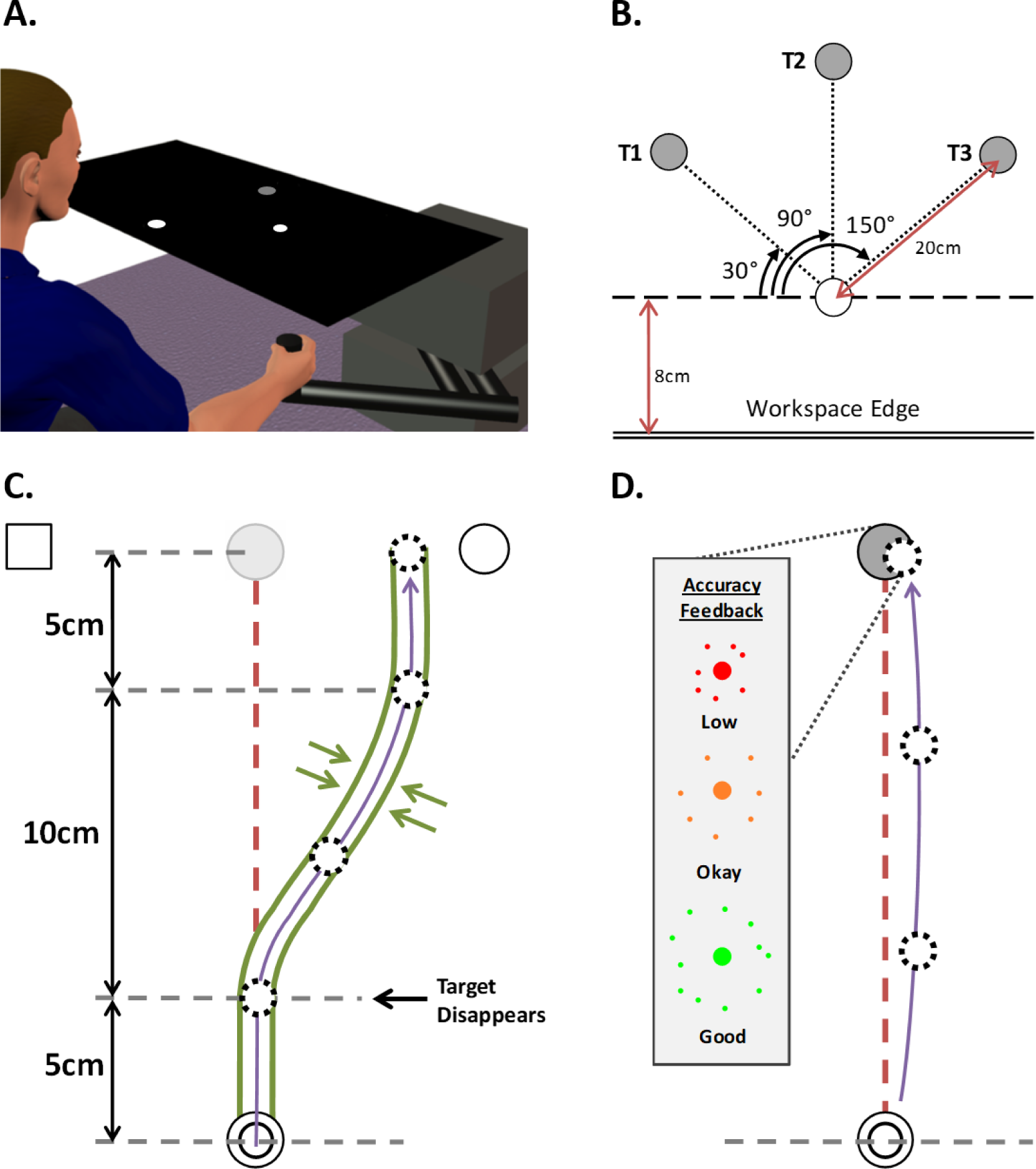
**A.** Example set-up of vBOT. LCD display (not shown) projects image onto mirrored surface to give visual feedback of hand location on robot handle. Mirror occludes any direct vision of the reaching arm **B.** Workspace locations and relative distances of the 3 targets (T1-T3) used in both the dynamic proprioception and rapid motor tasks **C.** Illustration of minimum jerk channel for the dynamic proprioception task. At termination, a circle and square are displayed to prompt a verbal response (“Circle” would be correct in this example). Target is visible for first 5cm before it disappears for remainder of trial, hand positon cursor remains occluded for all channel trials in a given block **D.** Illustration of rapid reaching task. Visual feedback of hand position was occluded once the cursor left the home position and remained so for the entire trial. Coloured feedback was provided at the target location on trial termination to indicate the endpoint accuracy of the movement. Both the experimental tasks in **C.** and **D.** are performed at target T2 (T1 and T3 not shown)

### Experimental Design

All participants performed the dynamic proprioceptive task first. Hence there was no possibility for the feedback associated with the rapid motor reaching task to alter or improve proprioceptive acuity to the same spatially located targets.

### Task 1: Dynamic Proprioception

#### Procedure

Participants made reaching movements towards 1 of the 3 targets with visual feedback of hand position occluded throughout, and target position occluded after the initial 5cm outward movement (see Figure 1C). These movements were constrained to a pre-defined minimum jerk path using stiff virtual walls (see Ostry et al. 2010) that steered the hand laterally away from the target (stiffness: 2000 N/m with 10 N.m/s damping imposed by vBOT motors; no force applied in the forward direction). At the end of the movement, the hand was held at the final deviated position and a white circle and square appeared at a constant position clockwise (CW) or counter-clockwise (CCW) of the target, respectively. The participant then verbally indicated the symbol (“Square” or “Circle”) which represented the side of the target they felt they had been guided to. With visual feedback of hand position still occluded, participants were actively guided back to the start position by a spring force (500 N/m, 1 N.m/s damping), where they remained until a new target appeared and the next trial began. The size of the lateral deviation was manipulated across trials by 2 randomly interleaved PEST sequences (see below).

#### PEST Sequences

The size and direction of the lateral deviation imposed by the virtual channels was dictated by two randomly interleaved PEST sequences (Taylor & Creelman, 1967) spanning across all 3 targets, with one starting each from the CCW (“Square”) and CW (“Circle”) sides of the target. In each block the initial deviation magnitude began at 3cm (±0.05cm added noise) with an initial step size of ±1cm, with 3 repeats (1 per target) at each “level” – the magnitude of the deviation. The deviation magnitude would increase or decrease depending on the cumulative accuracy of the 3 verbal responses per level. If participants made 2 or more correct responses, they would be deemed successful at that level and the deviation magnitude would reduce. However, if they scored 1 or fewer correct responses, the deviation magnitude would increase. Whenever the sequence reversed, the new step size was half of the previous one i.e. from 1cm to 0.5cm at the first reversal.

#### Outcome Measures and Analysis

The participant’s verbal responses were converted to binary values (“Circle” = 1, “Square” = 0) for each target; on the few occasions where there were multiple responses at the same deviation level to the same target, we then calculated the proportional response. A logistic function was then fitted to the data using the Matlab glmfit function to separately estimate the bias and uncertainly range of the psychometric response function. The bias represents the systematic or constant error in perception of hand position corresponding to the inverse of the 50^th^ percentile of the logistic function. Thus positive bias represents perception of hand position shifted towards the “Circle” (CW direction), and negative bias represents a perceptual shift towards the “Square” (CCW direction). The uncertainty range is defined as the interval between the 25^th^ and 75^th^ percentile of the fitted logistic function and represents a variable error in perception of hand position. To diminish the effects of outlying responses, data points which had a Pearson residual value which was more than 2 standard deviations away from the mean of the residuals were excluded from the analysis (this equated to roughly 4% of data).

Average movement speed was recorded for the portion of movement where the participant first reached 1cm from the start position to 1cm short of the final, deviated position. The mean orthogonal force imposed against the channel walls was also recorded in the middle of the final straight, 5cm portion of movement (16-19cm from the start; see Figure 1C). Both speed and lateral force were used as correlates for the bias to ensure that magnitude and direction of effort exerted against the channel wall was not influencing perceptual errors (Smith et al., 2009).

The dynamic proprioception task began with a short familiarisation block of 6 null-field and 9 perceptual channel trials. Participants then performed 5 blocks of 6 null-field trials followed by 48 channel trials with the opportunity for short breaks between blocks. The PEST sequence reset at the start of each new block such that the entire task included 5 PEST “runs” and totalled 80 perceptual judgements per target. Null-field trials were performed to the same spatially located targets and coloured feedback (an “explosion” graphic) was provided at the target location to indicate either a target “hit” or “miss”. These trials were intended to reduce proprioceptive drift during prolonged periods of occluded vision and were not analysed.

### Task 2: Rapid Motor Reaching

#### Procedure

Participants began each reaching trial by moving the visible hand position cursor to the start position. After a random wait time of between 2 and 3 seconds, one of the three targets appeared, and this was the participant’s cue to move towards the target as quickly and as accurately as possible. As soon as the cursor was moved outside of the start position it disappeared so the participant had no visual feedback of hand position during the movement. Participants were instructed to stop at their final position; the trial was terminated once hand velocity fell under 4cm/s at which point an animated “explosion” appeared at the target whose size and colour was based on the distance between the terminal hand position and the target (Figure 1D). Once the animation had finished, the hand position cursor reappeared, the target disappeared, and the participant was actively guided back towards the start position for the next trial.

#### Outcome Measures and Analysis

Kinematic performance was quantified by calculating reaction time (RT), peak hand velocity (PV), movement time (MT) and time to peak velocity (TPV). Movement initiation and termination were defined as the points where hand velocity first exceeded and then fell below 4cm/sec respectively. RT was therefore defined as the duration of time between the target appearing (i.e. movement initiation cue) and movement initiation. Trials where RT was less than 0.1sec or greater than 1sec were excluded from analysis (roughly 2% data). TPV was expressed as a percentage of total MT (time between movement initiation and termination) to examine the speed profile of the movement independently of its actual duration. Accuracy was quantified both by the absolute error (AE) at endpoint (the Euclidean distance from trial termination position to the target location) and by the lateral deviation at endpoint (LE). LE was calculated as the orthogonal distance from the linear path between start position and target, to endpoint and was included to improve the validity of the association with the proprioceptive measures, which also use an orthogonal deviation measure. Within participants variability in motor accuracy was assessed using the standard deviation of the accuracy measure across trials for each participant, separately for each target.

The rapid motor task was preceded by 9 practice trials (3 per target), with main task performance consisting of 3 blocks of 20 trials such that there were a total of 20 movements to each target.

### Physical Activity Measures

#### Older Adults

After completing the experiment, OAs were given wrist-worn accelerometers (Philips Actiwatch 2) to wear for 5 days (120 hours), where “activity counts” were logged in 30 second epochs. If an epoch had less than 40 counts it was deemed to be inactive (intermediate activity threshold defined by Philips Actiware software version 6.0.2). The sum of all counts in the surviving active epochs over the 5 days provided a physical activity (PA) metric for each older participant. The median value of the scores between participants was then used as a threshold to define “Inactive” and “Active” sub-groups of OAs for further analysis (demographic details for these groups are detailed in the Results section).

#### Younger Adults

We were unable to use accelerometer data to sub-group the YA participants. Hence self-reported PA measures were recorded for YAs using the IPAQ-Short questionnaire (Craig et al., 2003), with participants scoring in the highest “Health Enhancing Physical Activity” category being excluded from participation, in order to decrease heterogeneity.

### Working Memory

To test if working memory capacity influenced our proprioceptive measures, working memory was measured before participation in the experiment by using the backward digit span test, following previous reports of its use in proprioceptive ageing studies (Adamo et al., 2009; Goble et al., 2012). In this task, participants were required to memorise a sequence of random numbers (ranging 1-9; read out to them at a rate of approximately 1 number per second), and then recite them in reverse order. The task began with two trials at a sequence length of 2. If participants could correctly recite the sequence on at least 1 out of the 2 attempts at that sequence length level, the sequence length would increase by one. The task then incremented in this fashion until both attempted recitals were incorrect. The highest sequence length which the participant could correctly recite at least 1 out of the 2 attempts was recorded as their verbal working memory score.

### Statistical and Cross-Task Analysis

All data are presented as group means ± standard deviation unless otherwise stated, with values greater than 2.5 standard deviations away from the group mean at each target removed as outliers (approximately 5% of data). The remaining data were analysed in separate 3 × 3 mixed-design ANOVAs, with a between subjects factor of Group (inactive OAs, active OAs and YAs) and repeated measure of Target (T1-T3). A Greenhouse-Geisser correction was used in all cases where the sphericity assumption was violated, and significance was assessed at the α <.050 level. Statistically significant ANOVA effects and interactions were followed up with post-hoc t-test pairwise comparisons, and assessed for significance using a False Discovery Rate (FDR) analysis (Benjamini & Hochberg, 1995). The FDR analysis makes use of observed *p*-values to calculate an adjusted critical α-threshold, meaning it can be used in a range of different test statistics (Curran-Everett, 2000) as well as typically having higher power and being less conservative than other more commonly used methods, such as the Bonferroni correction (Benjamini & Hochberg, 1995). As such, it is gaining more popularity in the field of sensorimotor research (Boisgontier et al., 2014; Helsen et al., 2016). All *p*-values for multiple comparisons are therefore reported as uncorrected (Least Significant Difference) values but assessed at FDR adjusted α-thresholds (noted as α_FDR_). In situations where no comparisons are found to be significant, the smallest observed *p*-value (*p*_min_) and its associated critical significance threshold (still denoted as α_FDR_) is reported.

To assess the relationship between motor performance and proprioceptive acuity, a series of linear regression models were calculated. Since proprioceptive judgements were made along an axis orthogonal to the start-target vector, we assume that if either measure was related to motor control this would be most apparent with motor errors along a similar orthogonal axis. Thus, average lateral error (LE) and within-subject variation of LE (LE Var) were chosen as the motor performance measures to include in the regression models. Specifically, we hypothesize that proprioceptive noise could predict motor accuracy variation and so used uncertainty range to predict LE Var. We then examined the association between systematic proprioceptive and motor errors by using bias to predict LE. PA level was used as an additional predictor in the models which allowed us to collapse data across the inactive and active OA groups. Separate regression models were calculated for each of the 2 proprioceptive-motor relationships of interest for both OAs and YAs separately, with an FDR-adjusted α-threshold used to control for multiple tests.

## Results

### Physical Activity Grouping

The 31 OAs were divided into either a physically inactive or physically active sub-group according to a threshold median value of 1.68 × 10^6^ activity counts from the 5-day accelerometer data. This left 16 OAs in the inactive group (1.29 ±.31 × 10^6^counts; 7 male, 72.9 ± 5.1 yrs) and 15 in the active group (1.96 ±.26 × 10^6^counts; 4 male, 69.3 ± 2.7 yrs). The inactive group were found to be significantly older than the active group (t[22.9] = 2.5, *p* = .019); this difference is addressed directly as needed for cases where it could be deemed to have a confounding effect on pairwise comparisons.

### Dynamic Proprioception Task

#### Proprioceptive Measures

A summary of the proprioceptive outcome measures can be seen in Figures 2A (bias) and 2B (uncertainty range). There was a significant effect of Group on bias (F[2, 47] = 4.1, *p* = .023, *η*^*2*^_*p*_= .15) such that inactive OAs had larger biases than YAs (t[33] = 2.8, *p* = .009; α_FDR_= .017). Target also had a significant effect on bias (F[1.7, 78.6] = 3.8, *p* = .032, *η*^*2*^_*p*_= .08) but these differences did not survive FDR correction (*p*_min_ = .019; α_FDR_= .017). The interaction of Target × Group was not significant (F[3.3, 78.6] = .28, *p* = .861). To test whether the Group effect was truly due to physical inactivity of OAs and not their increased age (see Physical Activity Grouping) we correlated age and bias (averaged across all 3 targets) for the entire OA sample. The correlation was non-significant (r = .005, *p* = .977) and we conclude that the group effect on bias is indeed due to the physical inactivity of OAs.

Contrary to our predictions, there was no effect of Group on uncertainty range (F[2, 45] = .31, *p* = .733). There was an overall effect of Target (F[2, 90] = 4.8, *p* = .011, *η*^*2*^_*p*_ = .10), such that uncertainty range was larger at T3 than T2 (t[47] = −2.9, *p* = .006; α_FDR_= .017). There was no Group × Target interaction (F[4, 90] = .51, *p* = .730).

**Figure 2:**
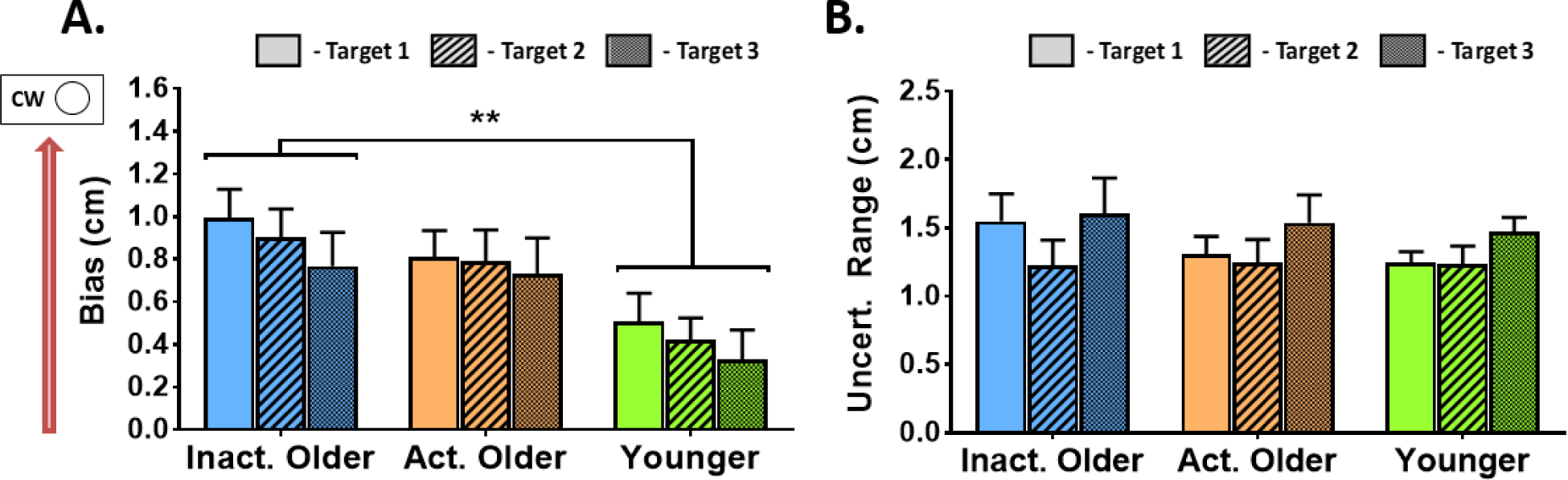
Group average data from dynamic proprioceptive task (mean ± standard error bars, effects of Target not shown) **A**. Results for bias, where inactive older adults had significantly larger, positive biases than younger adults (**\*\*** *p* <.010, multiple comparisons subjected to FDR adjusted α-threshold). Note all groups have positive biases which represents perception of hand position towards the clockwise (“Circle”) side of the targets **B**. Results for uncertainty range where there were no significant differences observed between any of the 3 groups

#### Kinematic Measures

Due to an unforeseen technical error, for 4 OAs in the physically inactive group we had only partial kinematic data which was non-analysable; the perceptual judgement data remained valid for all participants. For this reason kinematic data here was analysed as n = 12 for inactive OAs; the perceptual data for this sub-group did not differ from the others, tested with a mixed-ANOVA between the excluded and retained participants (bias *p* = .99, uncertainty range *p* = .16). YAs made the fastest movements (20.2 ± 5.9 cm/sec) followed by active OAs (16.1 ± 4.7 cm/sec) and inactive OAs who moved slowest (14.6 ± 5.4 cm/sec). Group had a significant effect on movement velocity (F[2, 43] = 4.6, *p* = .015, *η*^*2*^_*p*_ = .18) such that inactive OAs moved significantly slower than YAs (t[30] = −2.7, *p* = .012; α_FDR_= .017). Target also had a significant main effect on movement velocity (F[1.7, 71.7] = 18.3, *p* <.001, *η*^*2*^_*p*_ = .30), where pairwise comparisons revealed that movements were faster at T3 than both T1 (t[45] = −4.9, *p* <.001; α_FDR_ = .034) and T2 (t[45] = −4.5, *p* <.001). The Group × Target interaction was not significant (F[3.3, 71.7] = .73, *p* = .552).

Movement speed might influence perceptual performance in this task since the lateral acceleration through channel deviation (Figure 1C) would be greater for faster movements. We therefore tested if bias and uncertainty range were correlated with average movement velocity for each of the 3 different groups. We found that none of the correlations were significant for the bias (|r| <.34, *p*_min_ = .045; α_FDR_= .017); however the inactive OAs showed a significant, positive correlation between average movement velocity and uncertainty range (r = .46, *p* = .008; α_FDR_ = .017; all others |r| <.31) indicating faster movements were related to lower perceptual acuity. There were no significant relationships observed between bias and mean force exerted against the final section of the channel wall for any of the 3 groups (|r| <.294, *p*_min_ = .096; α_FDR_ = .017). This shows that systematic perceptual errors were independent of direction of effort exerted during the verbal reporting stage.

### Rapid Motor Reaching Performance

#### Performance Accuracy Measures

Results for the LE and LE Var motor accuracy measures are shown in Figure 3A and 3B respectively. All motor accuracy data (LE and AE parameters) are shown in Table 1.

**Table 1:** Group average motor performance accuracy measures for inactive older adults, active older adults, and younger adults. Values are given as means ± standard error, there were no significant group effects observed. LE = Lateral Endpoint Error, AE = Absolute Endpoint Error, in both cases Var = within-subject standard deviation (variation) in either measure

| Measure | Group | Target |  |  | <u>Overall</u> |
| --- | --- | --- | --- | --- | --- |
|  |  | 1 | 2 | 3 |  |
| **LE (cm)** | *Inactive Older* | .25 (± .13) | -1.24 (± .16) | -1.31 (± .20) | -.77 (± .12) |
|  | *Active Older* | .48 (± .15) | -1.42 (± .21) | -.97 (± .30) | -.64 (± .15) |
|  | *Younger* | -.40 (± .22) | -1.47 (± .17) | -1.08 (± .26) | -.98 (± .15) |
| **LE Var (cm)** | *Inactive Older* | .79 (± .06) | .86 (± .06) | .74 (± .05) | .80 (± .04) |
|  | *Active Older* | .94 (± .09) | .79 (± .07) | .95 (± .08) | .89 (± .07) |
|  | *Younger* | 1.03 (± .08) | .91 (± .05) | .99 (± .07) | .98 (± .05) |
| **AE (cm)** | *Inactive Older* | 1.57 (± .08) | 2.12 (± .19) | 2.03 (± .20) | 1.91 (± .13) |
|  | *Active Older* | 1.94 (± .13) | 2.34 (± .16) | 2.19 (± .20) | 2.16 (± .13) |
|  | *Younger* | 2.14 (± .11) | 2.24 (± .14) | 2.32 (± .18) | 2.23 (± .12) |
| **AE Var (cm)** | *Inactive Older* | .88 (± .06) | .94 (± .07) | .96 (± .07) | .92 (± .05) |
|  | *Active Older* | 1.06 (± .08) | .94 (± .07) | 1.09 (± .15) | 1.03 (± .09) |
|  | *Younger* | 1.08 (± .08) | 1.00 (± .06) | .99 (± .06) | 1.02 (± .06) |

**Figure 3:**
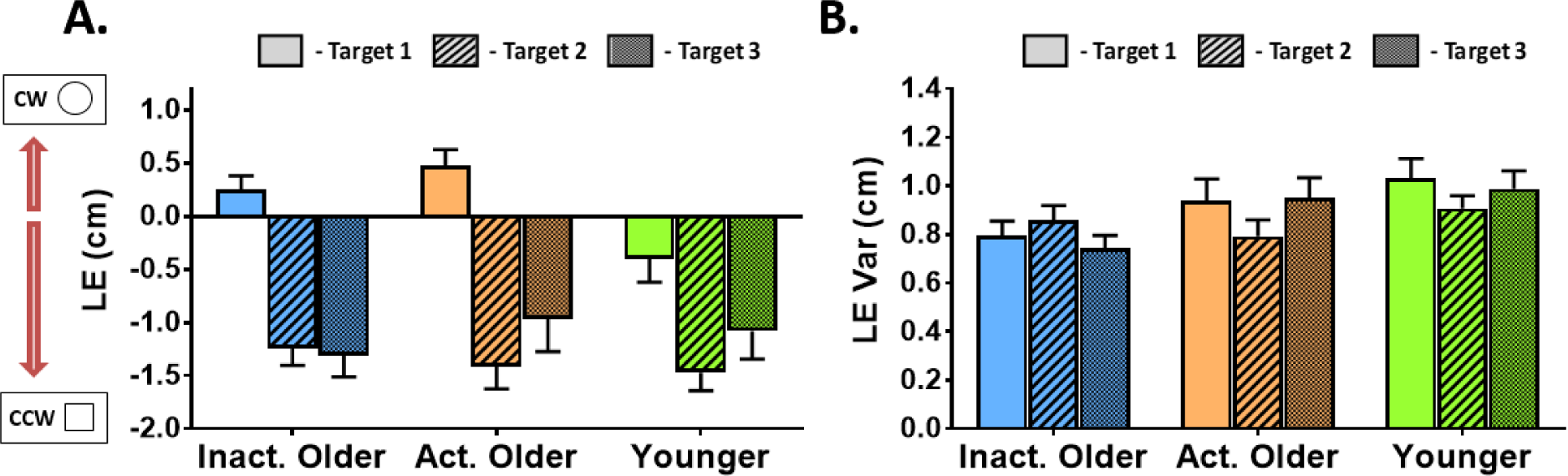
Group average motor performance accuracy measures (mean ± standard error bars) to be used in linear regression models with proprioceptive outcomes **A**. Results for lateral endpoint error (LE), where negative error represents an end-position which deviated laterally in the counter-clockwise (“Square” from the proprioceptive task) direction and vice versa **B**. Results for the within-subject variation (standard deviation) of the LE (LE Var). There were no significant differences between groups for either measure

The effect of Group on LE was not significant (F[2, 48] = 1.6, *p* = .218) but there was a significant effect of Target (F[1.4, 68.8] = 51.2, *p* <.001, *η*^*2*^_*p*_ = .52). Pairwise comparisons showed that LE was significantly different between all targets (T1 vs. T2, t[50] = 10.0, *p* <.001; T1 vs. T3, t[50] = 5.8, *p* <.001; T2 vs. T3, t[50] = −2.2, *p* = .035; α_FDR_ = .050), such that lateral errors were smallest at T1 and largest at T2. The interaction of Group and Target on LE was non-significant (F[2.9, 68.8] = 2.3, *p* = .091). There were no significant effects on LE Var for Group (F[2, 45] = 2.8, *p* = .072), Target (F[2, 90] = 1.2, *p* = .308) or their interaction (F[4, 90] = 1.8, *p* = .180). Thus, all groups had similar systematic and variable lateral endpoint errors in their movements.

There was also no effect of Group on AE (F[2, 44] = 1.8, *p* = .181) but there was a significant effect of Target (F[2, 88] = 7.6, *p* = .001, *η*^*2*^_*p*_ = .15) with endpoint errors being significantly larger at T2 (t[46] = −3.5, *p* = .001; α_FDR_ = .033) and T3 (t[46] = −2.7, *p* = .010) than at T1. The Group × Target interaction was non-significant (F[4, 88] = 1.11, *p* = .356). Neither Group (F[2, 44] = .78, *p* = .471) nor Target (F[1.7, 76.7) = .93, *p* = .389) had an effect on within-subject variation of AE (AE Var), with the interaction of Target × Group also being non-significant (F[3.5, 76.7] = 1.4, *p* = .260).

Collectively, this demonstrates a similar level of systematic and variable absolute errors between groups. This therefore shows endpoint accuracy in this motor task was maintained with advanced age, and was independent of PA.

Since participants were provided with accuracy feedback during the motor task, an additional ANOVA was performed on the accuracy measures in the early vs. late parts of the task (first vs. last 10 trials) to assess whether any motor learning occurred. We focus on, and report only, the factors of Time (early or late in the task) and Group × Time interaction effects from the 3 × 3 × 2 ANOVAs: (Group) × (Target) × (Time). There was a significant effect of Time on LE (F[1, 47] = 6.0, *p* = .018, *η*^*2*^_*p*_ = 0.11), AE (F[1, 42] = 6.2, *p* = .017, *η*^*2*^_*p*_ = .13) and AE Var (F[1, 42] = 7.0, *p* = .012, *η*^*2*^_*p*_ = .14) such that lateral errors, absolute errors and variation in absolute errors were all larger in the early stages of the task. However, there were no significant Group × Time interaction effects on any of the motor accuracy measures (all *p* >.050). This shows that although there were improvements in performance over the duration of the task, the extent of these improvements did not differ between the 3 groups.

#### Kinematic Performance Measures

The data for RT and PV are summarised in Figure 4A and 4B respectively, with all kinematic measures for the rapid motor task shown in Table 2. There was a significant effect of Group on RT (F[2, 47] = 11.5, *p* <.001, *η*^*2*^_*p*_ = .33) whereby both inactive OAs (t[19.7] = 4.6, *p* <.001; α_FDR_ = .033) and active OAs (t[18.1] = 3.7, *p* = .002) had longer reaction times than YAs. Likewise there was a significant effect of Target on RT (F[2, 94] = 15.0, *p* <.001, *η*^*2*^_*p*_ = .24) whereby participants reacted faster at target T1 compared to both T2 (t[49] = −4.1, *p* <.001; α_FDR_ = .033) and T3 (t[49] = −4.5, *p* <.001). The interaction effect of Group and Target on RT Group had a significant effect on PV (F[2, 46] = 18.8, *p* <.001, *η*^*2*^_*p*_ = .45), where both inactive OAs (t[33] = −5.2, *p* <.001; α_FDR_ = .033) and active OAs (t[32] = −4.5, *p* <.001) were significantly slower than YAs. Target also had a significant effect on PV (F[2, 92] = 32.8, *p* <.001, *η*^*2*^_*p*_ = .55), with pairwise comparisons showing each target was significantly different from one another (*p* ≤.001 in all cases; α_FDR_ = .050) such that T3 movements were fastest and T1 movements were slowest. The interaction effect of Group and Target on PV was also significant (F[4, 92] = 3.5, *p* = .011, *η*^*2*^_*p*_ = .13) with differences across targets most pronounced for the inactive OA group (Figure 4B). However, follow-up pairwise comparisons reflect the Group effect, in that both inactive and active OAs were significantly slower than YAs at all 3 targets (all *p* <.002; α_FDR_ = .033).

**Table 2:** Group average kinematic data (means ± standard error) for the rapid reaching task. Significant differences from younger adults are indicated by **\*\*** (*p* <.010) and **\*\*\*** (*p* <.001; multiple comparisons subjected to FDR adjusted α-threshold). React. Time = Reaction Time, Peak Vel. = Peak Hand Velocity, Move. Time = Movement Time, TPV = Time to Peak Velocity

| Measure | Group | Target |  |  | <u>Overall</u> |
| --- | --- | --- | --- | --- | --- |
|  |  | 1 | 2 | 3 |  |
| **React. Time (sec)** | *Inactive Older* | .44 (± .02) | .46 (± .02) | .47 (± .02) | ***.46 (± .02) |
| | *Active Older* | .42 (± .02) | .46 (± .02) | .46 (± .03) | **.45 (± .02) |
| | *Younger* | .35 (± .01) | .37 (± .01) | .36 (± .01) | .36 (± .01) |
| **Peak Vel. (cm/sec)** | *Inactive Older* | ***48.1 (± 3.2) | ***51.2 (± 3.9) | **57.6 (± 3.9) | ***52.3 (± 3.6) |
| | *Active Older* | ***53.0 (± 3.7) | ***56.8 (± 3.5) | **58.1 (± 4.1) | ***55.9 (± 3.7) |
| | *Younger* | 81.4 (± 4.0) | 84.3 (± 4.8) | 85.3 (± 4.7) | 83.7 (± 4.5) |
| **Move. Time (sec)** | *Inactive Older* | ***.77 (± .05) | ***.73 (± .05) | ***.65 (± .04) | ***.72 (± .05) |
| | *Active Older* | ***.73 (± .05) | ***.68 (± .04) | ***.66 (± .04) | ***.69 (± .04) |
| | *Younger* | .49 (± .02) | .46 (± .02) | .45 (± .01) | .47 (± .02) |
| **TPV (% Move Duration)** | *Inactive Older* | 42.5 (± 1.2) | 43.6 (± 1.3) | 47.4 (± 1.5) | 44.5 (± 1.2) |
| | *Active Older* | 41.1 (± 1.3) | 42.7 (± 1.7) | 44.6 (± 1.5) | 42.8 (± 1.4) |
| | *Younger* | 42.9 (± .5) | 44.2 (± .6) | 46.0 (± .6) | 44.4 (± .5) |

**Figure 4:**
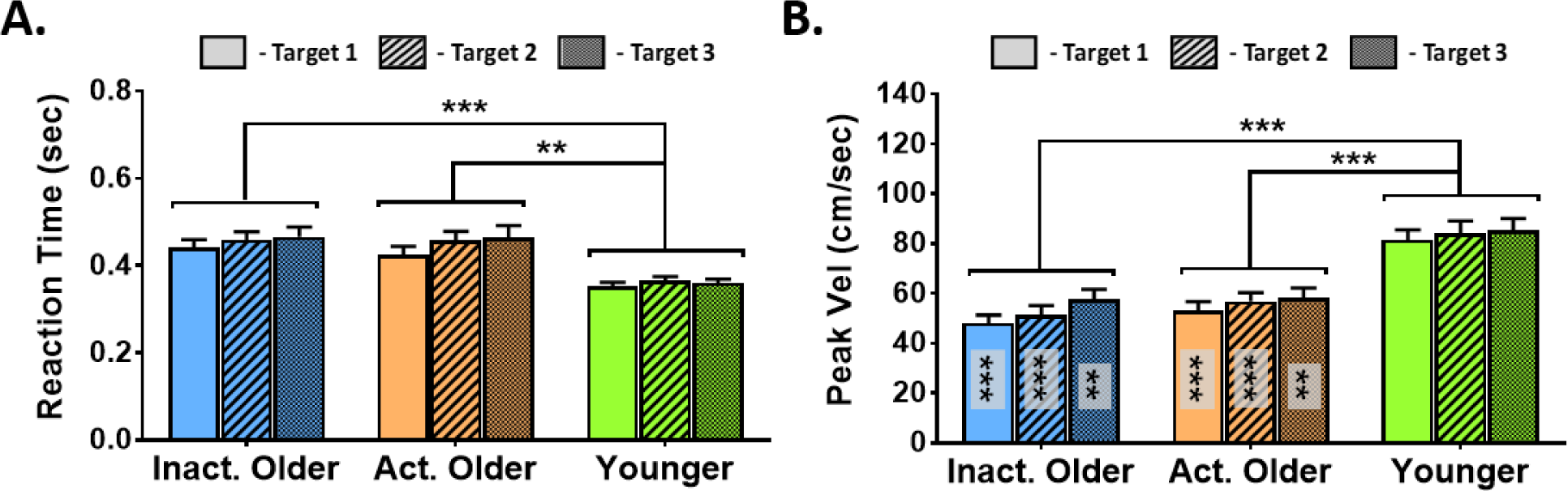
Group average kinematic data (mean ± standard error bars) for reaction time (**A**) and peak hand velocity (**B**) in the rapid reaching task. Significant differences from younger adults are indicated by **\*\*** (*p* <.010), **\*\*\*** (*p* <.001) in the upper section, with multiple comparisons subjected to FDR adjusted α-threshold. Asterisks within bars in panel B denote significant differences from younger adults at same target

There was a significant effect of Group on MT (F[2, 47] = 15.0, *p* <.001, *η*^*2*^ = .39), such that both inactive OAs (t[18.0] = 4.9, *p* <.001; α_FDR_ = .033) and active OAs (t[17.5] = 4.8, *p* <.001) made longer duration movements than YAs. There was also a main effect of Target (F[1.5, 72.3] = 45.3, *p* <.001, *η*^*2*^_*p*_ = .49) where all targets were significantly different from one another (all *p* <.001; α_FDR_ = .050) such that movements were made with the shortest duration to T3 and longest to T1. The Group by Target interaction was also significant for MT (F[3.1, 72.3] = 5.2, *p* = .003, *η*^*2*^_*p*_ = .18), but as with the peak velocity measure, follow-up pairwise comparisons reflected the main effect of Group with both inactive (all *p* <.001; α_FDR_ = .033) and active OAs (all *p* <.001) displaying longer movement durations than YAs at all targets.

The main effect of Group on TPV was not significant (F[2, 47] = .77, *p* = .473). However, there was a main effect of Target (F[2, 94] = 33.7, *p* <.001, *η*^*2*^_*p*_ = .42) whereby TPV was significantly different between all 3 targets (*p* <.002 in all cases; α_FDR_ = .050) such that peak velocity occurred later in movements to T3 and earlier in movements to T1. There was no interaction of Group and Target on DPV (F[4, 94] = 1.1, *p* = .382).

Together, the results from these kinematic measures shows that there were target-specific common kinematic features across all three groups, but overall, the OAs tend to react and move more slowly than YAs, regardless of their PA level. However, the shape of velocity profiles of movements were similar between all groups.

#### Speed-Accuracy Trade-off

Since there were significant differences in peak hand velocity between older and younger groups, we wanted to test for a potential speed-accuracy trade-off. We therefore divided both LE and AE values by corresponding PV on a trial-by-trial basis to create lateral and absolute error indices controlled for movement speed (LE_PVCont_ and AE_PVCont_ respectively), then analysed by 3 × 3 mixed-design ANOVAs: (Group) × (Target), as above.

There was no effect of Group on LE_PVCont_ (F[2, 46] = .19, *p* = .826) but the main effect of Target was significant (F[1.6, 73.7] = 58.1, *p* <.001, *η*^*2*^_*p*_ = .56; see Figure 5A). Pairwise comparisons showed that velocity controlled lateral errors were significantly different between all targets (T1 vs. T2, t[48] = 9.2, *p* <.001; T1 vs. T3, t[48] = 6.0, *p* <.001; T2 vs. T3, t[48] = −2.4, *p* = .018; α_FDR_ = .050), with smallest errors at T1 and largest at T2. The Group × Target interaction LE_PVCont_ was also significant (F[3.2, 73.7] = 4.8, *p* = .004, *η*^*2*^_*p*_ = .17). There was a trend towards both active (t[33] = 2.6, *p* = .0063, α_FDR_ = .0056) and inactive (t[33] = 2.6, *p* = .015) OAs having more positive velocity controlled lateral errors than YAs at T1, but these effects did not survive FDR correction (*p*_min_ = .085 for other of 3 [Group] × 3 [Target] comparisons).

**Figure 5:**
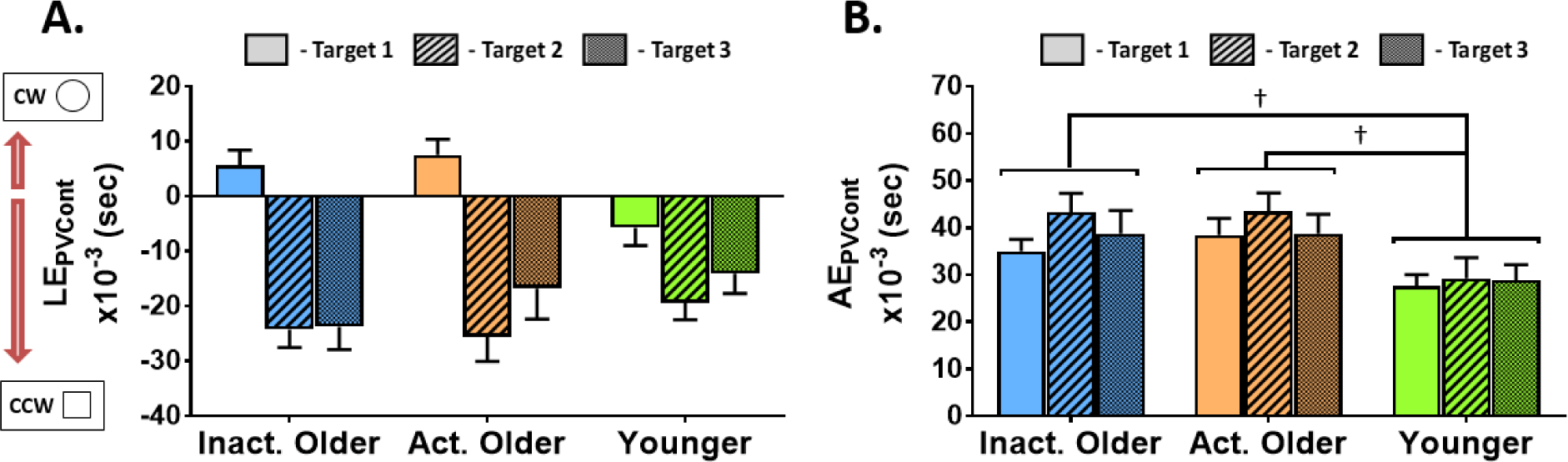
Group average motor accuracy measures controlled for by peak hand velocity (means ± standard error). **A.** Lateral error divided by peak hand velocity (LE_PVCont_) where more positive values represent errors to the clockwise (or “Circle” from proprioceptive task) side. **B.** Absolute errors divided by peak hand velocity (AE_PVCont_). Pairwise comparisons which were significant (*p* <.05) but did not survive corrections for multiple comparisons are indicated by **†**

The Group effect on AE_PVCont_ was significant (F[2, 42] = 4.2, *p* = .021, *η*^*2*^_*p*_ = .17; Figure 5B) but follow-up pairwise comparisons did not reveal any specific group differences after FDR correction, despite both active (t[30] = 2.5, *p* = .0171; α_FDR_ = .0166) and inactive (t[30] = 2.2, *p* = .035) OAs showing trends towards having larger velocity controlled absolute errors than YAs. There was also a significant main effect of Target (F[2, 84] = 4.2, *p* = .023, *η*^*2*^_*p*_ = .09) but follow-up pairwise comparisons were not significant following FDR correction (*p*_min_ = .020; α_FDR_ = .017). The Group × Target interaction on AE_PVCont_ was not significant (F[4, 84] = .73, *p* = .574).

Collectively, this additional analysis of the speed-accuracy trade-off shows that the maintenance of absolute endpoint accuracy in OAs may be partially explained by movement slowing. However, the lateral errors appear to be similar between age groups even when controlling for movement speed, suggesting they may be less susceptible to a speed-accuracy trade-off in this context.

### Working Memory Capacity

All groups had similar working memory capacity scores, as indicated by a non-significant one-way ANOVA (F[2, 48] = .16, *p* = .854). YAs had the highest score (5.8 ± 1.6 numbers recalled) followed by active OAs (5.7 ± 1.4) and inactive OAs with the lowest score (5.5 ± 1.3). To test if working memory was related to proprioceptive performance, we correlated the bias and uncertainty range, averaged across all 3 targets, with working memory score. There were no significant relationships found (all |r| <.38, *p*_min_ = .106; α_FDR_ = .008), showing proprioceptive performance was independent of working memory.

### Predicting Motor Performance from Proprioceptive Acuity

To allow visual comparison of the reaching performance with the proprioceptive measures, the spatial distribution of individuals’ average end-positions and the 95% confidence interval ellipses in the motor reaching task are shown in Figure 6 for each target, with the bias and uncertainty range from the proprioceptive task shown in bar-format.

**Figure 6:**
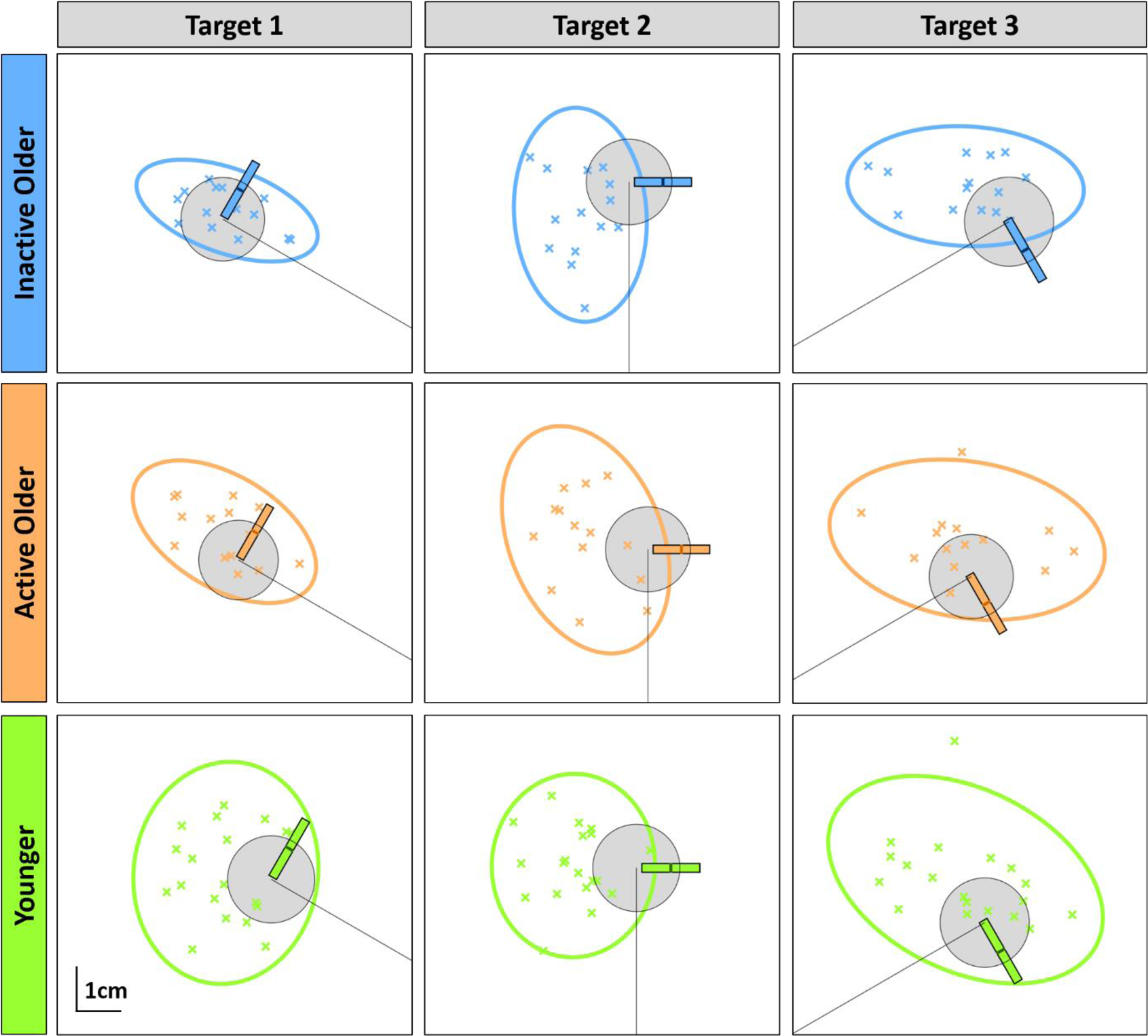
Individual participant average end-positions from rapid motor task (coloured ‘X’ markers) and 95% confidence ellipses for each of the different groups and targets. Group average data from dynamic proprioceptive task is scaled and superimposed over targets as coloured bars. The central thick coloured line in each bar represents the bias and on average shows participants perceived their hand to be more towards the clockwise (“Circle”) side of the target. The length of the coloured bar represents the uncertainty range and was similar between groups (figure generated for visualisation purposes only)

We generated 2 regression models for each proprioceptive-motor performance pairing, collapsing data across all 3 targets, giving 4 models overall. Neither the bias and LE (OAs, R^2^ = .002; YAs, R^2^ = .020) nor the uncertainty range and LE Var (OAs, R^2^ = .060; YAs, R^2^= .035; *p*_min_= .090; α_FDR_ = .013) models were significant (see Table 3 for summary). We did observe that uncertainty range was a significant, negative predictor of LE Var for OAs only (β = −.245; *p* = .030), however, this did not survive corrections for multiple comparisons and the overall model still accounted for only 6% of the variance in the data. The lack of relationship between proprioceptive uncertainty and motor error in advanced age contradicts our original prediction, and no consistent positive association was seen in any group.

**Table 3:** Summary of statistics for linear regression models predicting motor accuracy from proprioceptive and physical activity (PA) measures. Upper panel shows lateral error (LE) predicted by bias and PA, lower panel shows lateral error variability (LE Var) predicted by uncertainty range (UncR) and PA. All models were non-significant (*p*_min_= .090; α_FDR_= .013), with **†** indicating significant standardized coefficient (*p* <.05) which did not survive corrections for multiple comparisons.

| Model | Group | Model Measure |  |  |
| --- | --- | --- | --- | --- |
|  |  | R<sup>2</sup> | Propriocept.<br>β-Coeff. | PA β-Coeff. |
| **LE predicted by Bias and PA** | *Older* | .002 | -.022 | .041 |
|  | *Younger* | .020 | -.137 | .022 |
| **LE Var predicted by UncR and PA** | *Older* | .060 | -.245<sup>†</sup> | .002 |
|  | *Younger* | .035 | -.152 | -.103 |

## Discussion

This experiment aimed to determine the relationship between dynamic proprioceptive acuity and movement control in the upper limb with advanced age. Although we found stereotypical features of ageing in motor kinematics, we also found that proprioceptive bias, and not uncertainty range, was larger for physically inactive OAs, contrasting to our predictions. While we did observe a trend towards higher uncertainty range predicting lower variability in motor accuracy for OAs, the direction of this relationship and its limited strength (R^2^ = .06) lead us to conclude a negligible association overall. Ultimately, proprioceptive uncertainty was not consistently related to variability in movement accuracy; thus, we find no evidence to link proprioception and movement control in either older or younger adults in this experiment.

Our results replicate the findings of Helsen et al. (2016), who showed a dissociation of proprioceptive acuity and rapid motor performance, but we extend beyond their results to show this is true when proprioception is measured via an active movement task, which more closely mimics the sensation involved in voluntary movement. Helsen et al. (2016) concluded that OAs were able to overcome a decline in sensory acuity through increased reliance on predictive control mechanisms in a “play-it-safe” strategy (Elliott et al., 2010). We also saw evidence that OAs tend to emphasise accuracy over speed, exemplified by their increased reaction times and reduced peak velocities. These speed differences may partially explain the comparable endpoint accuracy seen between groups (Figure 5B); a finding which has also been reported elsewhere (Helsen et al., 2016; Lee et al., 2007; Seidler-Dobrin & Stelmach, 1998). We note that the utility of online proprioceptive feedback in fast, discrete, movements is likely reduced compared to slower, guided movements, and the reliance on predictive mechanisms may therefore already be high in our reaching task (Miall & Wolpert, 1996; Shadmehr, Smith, & Krakauer, 2010; Wolpert, Ghahramani, & Jordan, 1995). However, if OAs do tend to favour accuracy over speed, as our data suggest and as others have argued (Forstmann et al., 2011), then it seems unlikely they would opt to make movements so rapidly that feedback control would be completely negated. In the future, it may therefore be interesting to examine the relationship between proprioception and motor control in movement tasks that deliberately emphasise sensory guidance. This could include more continuous movements such as circular tracking (Levy-Tzedek, 2017), in which OAs increase movement radius and speed to a greater extent than YAs, upon removal of visual feedback. Alternatively, training in the control of objects in virtual environments, such as the ball balancing task reported recently by Elangovan, Cappello, Masia, Aman, & Konczak (2017), which increases proprioceptive acuity of the wrist. But perhaps a more commonly employed paradigm that can probe proprioceptive regulation of motor control is adaptation to novel field dynamics, where mechanical perturbations to the arm create unexpected trajectory deviations during reaching (Shadmehr & Mussa-Ivaldi, 1994). In ageing, this task has been studied surprisingly scarcely, with mixed findings on the extent to which adaptation is impaired in later life (Cesqui, Macri, Dario, & Micera, 2008; Huang & Ahmed, 2014; Reuter, Pearcey, & Carroll, 2018; Trewartha, Garcia, Wolpert, & Flanagan, 2014). Considering proprioceptive feedback is necessary to minimise within-trial performance errors in these tasks (Miall et al., 2018; Sarlegna, Malfait, Bringoux, Bourdin, & Vercher, 2010; Yousif, Cole, Rothwell, & Diedrichsen, 2015), it may be that proprioceptive acuity could account for some of the reported variance in age-related adaptation impairments. Moreover, since we report age and physical activity effects on proprioceptive bias, it would be interesting to see whether older participants recalibrate their proprioceptive sensation with forcefield learning in a similar way to YAs (Ostry et al., 2010), and if this predicts their adaptive performance.

Contrary to our predictions and to prior literature, we showed that physical inactivity did not increase proprioceptive uncertainty in OAs. We suggest this novel finding reflects the steps we took to remove confounds when measuring proprioception. Namely, we used active instead of passive movements (Smith et al., 2009) which minimises position matching errors in both older and younger adults (Erickson & Karduna, 2012; Langan, 2014; Lönn et al., 2000). We also required instantaneous perceptual judgements to minimise age-dependent memory effects (Goble et al., 2012), and we avoided comparison between the two arms to minimise effects of central degeneration, which may compromise interhemispheric transfer of sensorimotor signals (Hou & Pakkenberg, 2012). Instead, we were able to measure a significant increase in systematic perceptual error for the physically inactive OAs. Proprioceptive biases have been well established for reaching and pointing movements (Cressman et al., 2010; van Beers, Sittig, & van der Gon, 1998; Vindras, Desmurget, Prablanc, & Viviani, 1998; Wilson, Wong, & Gribble, 2010) and perception of limb position is frequently biased towards the side of the body where the limb is tested. These biases have been shown to be dependent on several task-specific factors, such as reach distance (van Beers et al., 1998; Wilson et al., 2010), limb used (Wilson et al., 2010; Wong, Wilson, Kistemaker, & Gribble, 2014) and whether visual or haptic reference positions are used (Kuling, Brenner, & Smeets, 2016). Less is known about individual differences which influence the presentation of these errors, or the mechanism by which they may occur. Here, we have shown that physical inactivity in ageing is a contributing factor. Although the cause is as yet unclear, a reduction in physical activity could lead to everyday limb movements being made within a more concentrated volume, ipsilateral to the limb (Howard, Ingram, Körding, & Wolpert, 2009), biasing sensory experience to this region. Increased sensory uncertainty upon removal of vision (as in the proprioceptive assessment task) may therefore lead to greater reliance on prior experience during the optimal estimation of limb position (Gritsenko, Krouchev, & Kalaska, 2007; Körding & Wolpert, 2006). We also note that spindle afferents are directionally tuned to specific movements (Bergenheim, Ribot-Ciscar, & Roll, 2000; Jones, Wessberg, & Vallbo, 2001) and loss of intrafusal fibres with age has been shown to be muscle specific (Kararizou, Manta, Kalfakis, & Vassilopoulos, 2005). Therefore if movements are indeed limited to a smaller range in physically inactive adults, a selective loss of intrafusal fibres which are directionally tuned to the less frequent movements, might result. Collectively, these effects could lead to the increase in proprioceptive bias we observed in the physically inactive OAs.

Unfortunately, the wrist-worn accelerometers we used do not provide spatial information, and this suggestion remains to be tested. An alternative could be that the perceptual bias arose from proprioceptive drift (Brown, Rosenbaum, & Sainburg, 2003b, 2003a; Desmurget, Vindras, Gréa, Viviani, & Grafton, 2000). However, drift is typically observed during repetitive, unconstrained movements and has been attributed to the persistence of motor errors rather than to proprioceptive fading (Brown et al., 2003b). In addition, the extent of proprioceptive drift has been associated with movement speed (Brown et al., 2003b), and we found no association between bias and movement velocity.

We do, however, report a positive correlation of average movement speed and uncertainty range in the proprioceptive task for the inactive OAs. This observation may further reflect a speed-accuracy trade-off where insufficient sensory information is accumulated to make reliable perceptual judgements as movement speed increases (Bogacz, Wagenmakers, Forstmann, & Nieuwenhuis, 2010; Heekeren, Marrett, & Ungerleider, 2008). In advanced age there is a high susceptibility to prefrontal cortex degeneration (Giorgio et al., 2010; Salat, 2004) which can be mediated by physical inactivity (Colcombe et al., 2003). Both attention and memory depend on these frontal brain regions and have been reported to influence the accuracy of limb position matching (Goble et al., 2012). Limited cognitive resources in the inactive OAs might therefore impair their ability to process sensory feedback for perceptual judgements. However, we found no relationship between verbal working memory score and perceptual acuity for any group, suggesting this is not a factor in our inactive elderly group.

In conclusion, we found systematic differences in movement kinematics in OAs compared to YAs, as expected from previous reports. We also found an age-dependent increase in proprioceptive bias measured in active, multi-joint movement, but not of uncertainty range. This finding is novel and may reflect our careful task design which aimed to remove methodological confounds for testing with an ageing population. However, we did not find any evidence to suggest that proprioceptive acuity is related to performance in rapid, goal-orientated movement, in either older or younger adults. The relationship between proprioceptive acuity and motor control remains uncertain, and warrants further investigation under movement conditions which emphasise the utility of online proprioceptive feedback.

## Acknowledgements

This work was funded by the MRC-ARUK Centre for Musculoskeletal Ageing Research (CMAR) and by the Wellcome Grant WT087554.

## References

1. Adamo, D. E., Alexander, N. B., & Brown, S. H. (2009). The Influence of Age and Physical Activity on Upper Limb Proprioceptive Ability. J Aging Phys Act, 17(3), 272–293.

2. Adamo, D. E., Martin, B. J., & Brown, S. H. (2007). Age-related differences in upper limb proprioceptive acuity. Perceptual and Motor Skills, 104(3 Pt 2), 1297–1309.

3. Benjamini, Y., & Hochberg, Y. (1995). Controlling the false discovery rate: a practical and powerful approach to multiple testing. Journal of the Royal Statistical Society. Series B (Methodological), 289–300.

4. Bergenheim, M., Ribot-Ciscar, E., & Roll, J.-P. (2000). Proprioceptive population coding of two-dimensional limb movements in humans: I. Muscle spindle feedback during spatially oriented movements. Experimental Brain Research, 134(3), 301–310.

5. Bogacz, R., Wagenmakers, E.-J., Forstmann, B. U., & Nieuwenhuis, S. (2010). The neural basis of the speed–accuracy tradeoff. Trends in Neurosciences, 33(1), 10–16.

6. Boisgontier, M. P., Olivier, I., Chenu, O., & Nougier, V. (2012). Presbypropria: the effects of physiological ageing on proprioceptive control. AGE, 34(5), 1179–1194.

7. Boisgontier, M. P., Van Halewyck, F., Corporaal, S. H. A., Willacker, L., Van Den Bergh, V., Beets, I. A. M.,…Swinnen, S. P. (2014). Vision of the active limb impairs bimanual motor tracking in young and older adults. Frontiers in Aging Neuroscience, 6.

8. Brown, L. E., Rosenbaum, D. A., & Sainburg, R. L. (2003a). Limb Position Drift: Implications for Control of Posture and Movement. Journal of Neurophysiology, 90(5), 3105–3118.

9. Brown, L. E., Rosenbaum, D. A., & Sainburg, R. L. (2003b). Movement speed effects on limb position drift. Experimental Brain Research, 153(2), 266–274.

10. Ceballos, D., Cuadras, J., Verdu, E., & Navarro, X. (1999). Morphometric and ultrastructural changes with ageing in mouse peripheral nerve. Journal of Anatomy, 195(4), 563–576.

11. Cesqui, B., Macri, G., Dario, P., & Micera, S. (2008). Characterization of age-related modifications of upper limb motor control strategies in a new dynamic environment. Journal of NeuroEngineering and Rehabilitation, 5(1), 31.

12. Colcombe, S. J., Erickson, K. I., Raz, N., Webb, A. G., Cohen, N. J., McAuley, E., & Kramer, A. F. (2003). Aerobic Fitness Reduces Brain Tissue Loss in Aging Humans. The Journals of Gerontology Series A: Biological Sciences and Medical Sciences, 58(2), M176–M180.

13. Contreras-Vidal, J. L., Teulings, H. L., & Stelmach, G. E. (1998). Elderly Subjects are Impaired in Spatial Coordination of Fine Motor Control. Acta Psychologica, 100, 25–35.

14. Craig, C. L., Marshall, A. L., Sjöström, M., Bauman, A. E., Booth, M. L., Ainsworth, B. E.,…Oja, P. (2003). International Physical Activity Questionnaire: 12-Country Reliability and Validity: Medicine & Science in Sports & Exercise, 35(8), 1381–1395.

15. Cressman, E. K., & Henriques, D. Y. P. (2009). Sensory Recalibration of Hand Position Following Visuomotor Adaptation. Journal of Neurophysiology, 102(6), 3505–3518.

16. Cressman, E. K., Salomonczyk, D., & Henriques, D. Y. P. (2010). Visuomotor adaptation and proprioceptive recalibration in older adults. Experimental Brain Research, 205(4), 533–544.

17. Curran-Everett, D. (2000). Multiple comparisons: philosophies and illustrations. American Journal of Physiology-Regulatory, Integrative and Comparative Physiology, 279(1), R1–R8.

18. Darling, W. G., Cooke, J. D., & Brown, S. H. (1989). Control of simple arm movements in elderly humans. Neurobiology of Aging, 10(2), 149–157.

19. Desmurget, M., Vindras, P., Gréa, H., Viviani, P., & Grafton, S. T. (2000). Proprioception does not quickly drift during visual occlusion. Experimental Brain Research, 134(3), 363–377.

20. Elangovan, N., Cappello, L., Masia, L., Aman, J., & Konczak, J. (2017). A robot-aided visuo-motor training that improves proprioception and spatial accuracy of untrained movement. Scientific Reports, 7(1).

21. Elliott, D., Hansen, S., Grierson, L. E. M., Lyons, J., Bennett, S. J., & Hayes, S. J. (2010). Goal-directed aiming: Two components but multiple processes. Psychological Bulletin, 136(6), 1023–1044.

22. Erickson, R. I. C., & Karduna, A. R. (2012). Three-dimensional repositioning tasks show differences in joint position sense between active and passive shoulder motion. Journal of Orthopaedic Research, 30(5), 787–792.

23. Forstmann, B. U., Tittgemeyer, M., Wagenmakers, E.-J., Derrfuss, J., Imperati, D., & Brown, S. (2011). The Speed-Accuracy Tradeoff in the Elderly Brain: A Structural Model-Based Approach. Journal of Neuroscience, 31(47), 17242–17249.

24. Giorgio, A., Santelli, L., Tomassini, V., Bosnell, R., Smith, S., De Stefano, N., & Johansen-Berg, H. (2010). Age-related changes in grey and white matter structure throughout adulthood. NeuroImage, 51(3), 943–951.

25. Goble, D. J., Coxon, J. P., Wenderoth, N., Van Impe, A., & Swinnen, S. P. (2009). Proprioceptive sensibility in the elderly: Degeneration, functional consequences and plastic-adaptive processes. Neuroscience & Biobehavioral Reviews, 33(3), 271–278.

26. Goble, D. J., Mousigian, M. A., & Brown, S. H. (2012). Compromised encoding of proprioceptively determined joint angles in older adults: the role of working memory and attentional load. Experimental Brain Research, 216(1), 35–40.

27. Gritsenko, V., Krouchev, N. I., & Kalaska, J. F. (2007). Afferent Input, Efference Copy, Signal Noise, and Biases in Perception of Joint Angle During Active Versus Passive Elbow Movements. Journal of Neurophysiology, 98(3), 1140–1154.

28. Heekeren, H. R., Marrett, S., & Ungerleider, L. G. (2008). The neural systems that mediate human perceptual decision making. Nature Reviews Neuroscience, 9(6), 467–479.

29. Helsen, W. F., Van Halewyck, F., Levin, O., Boisgontier, M. P., Lavrysen, A., & Elliott, D. (2016). Manual aiming in healthy aging: does proprioceptive acuity make the difference? Age (Dordr), 38(2).

30. Herter, T. M., Scott, S. H., & Dukelow, S. P. (2014). Systematic Changes in Position Sense Accompany Normal Aging Across Adulthood. Journal of Engineering and Rehabilitation, 11(43).

31. Hou, J., & Pakkenberg, B. (2012). Age-related degeneration of corpus callosum in the 90+ years measured with stereology. Neurobiology of Aging, 33(5), 1009.e1–1009.e9.

32. Howard, I. S., Ingram, J. N., Körding, K. P., & Wolpert, D. M. (2009). Statistics of Natural Movements Are Reflected in Motor Errors. Journal of Neurophysiology, 102(3), 1902–1910.

33. Howard, I. S., Ingram, J. N., & Wolpert, D. M. (2009). A modular planar robotic manipulandum with end-point torque control. Journal of Neuroscience Methods, 181(2), 199–211.

34. Huang, H. J., & Ahmed, A. A. (2014). Older adults learn less, but still reduce metabolic cost, during motor adaptation. Journal of Neurophysiology, 111(1), 135–144.

35. Hurley, M. V., Rees, J., & Newham, D. J. (1998). Quadriceps Function, Proprioceptive Acuity and Functional Performance in Healthy Young, Middle-Aged and Elderly Subjects. Age and Ageing, 27, 55–62.

36. Jacobs, J. M., & Love, S. (1985). Qualitative and quantitative morphology of human sural nerve at different ages. Brain, 108(4), 897–924.

37. Jones, K. E., Wessberg, J., & Vallbo, Å. B. (2001). Directional Tuning of Human Forearm Muscle Afferents During Voluntary Wrist Movements. Journal of Physiology, 536.2, 635–647.

38. Kararizou, E., Manta, P., Kalfakis, N., & Vassilopoulos, D. (2005). Morphometric Study of the Human Muscle Spindle. Anal Quant Cytol Histol, 27(1), 1–4.

39. Ketcham, C. J., Seidler, R. D., Van Gemmert, A. W. A., & Stelmach, G. E. (2002). Age-Related Kinematic Differences as Influenced by Task Difficulty, Target Size, and Movement Amplitude. J Gerontol B Psychol Sci Soc Sci, 57, 54–64.

40. Körding, K. P., & Wolpert, D. M. (2006). Bayesian decision theory in sensorimotor control. Trends in Cognitive Sciences, 10(7), 319–326.

41. Kuling, I. A., Brenner, E., & Smeets, J. B. J. (2016). Errors in visuo-haptic and haptic-haptic location matching are stable over long periods of time. Acta Psychologica, 166, 31–36.

42. Langan, J. (2014). Older adults demonstrate greater accuracy in joint position matching using self-guided movements. Human Movement Science, 36, 97–106.

43. Lee, G., Fradet, L., Ketcham, C. J., & Dounskaia, N. (2007). Efficient control of arm movements in advanced age. Experimental Brain Research, 177(1), 78–94.

44. Lei, Y., & Wang, J. (2018). The effect of proprioceptive acuity variability on motor adaptation in older adults. Experimental Brain Research, 236(2), 599–608.

45. Levy-Tzedek, S. (2017). Motor errors lead to enhanced performance in older adults. Scientific Reports, 7(1).

46. Lexell, J. (1995). Human aging, muscle mass, and fiber type composition. The Journals of Gerontology. Series A, Biological Sciences and Medical Sciences, 50 Spec No, 11–16.

47. Lönn, J., Crenshaw, A. G., Djupsjöbacka, M., Pedersen, J., & Johansson, H. (2000). Position Sense Testing. Influence of Starting Position and Type of Displacement. Archives of Physical Medicine and Rehabilitation, 81(5), 592–597.

48. Lord, S. R., Clark, R. D., & Webster, I. W. (1991). Postural Stability and Associated Physiological Factors in a Population of Aged Persons. Journal of Gerontology, 46(3), M69–76.

49. Miall, R. C., Kitchen, N. M., Nam, S.-H., Lefumat, H., Renault, A. G., Ørstavik, K.,…Sarlegna, F. R. (2018). Proprioceptive loss and the perception, control and learning of arm movements in humans: evidence from sensory neuronopathy. Experimental Brain Research, 236(8), 2137–2155.

50. Miall, R. C., & Wolpert, D. M. (1996). Forward Models for Physiological Motor Control. Neural Networks, 9(8), 1265–1279.

51. Morley, J. E., Baumgartner, R. N., Roubenoff, R., Mayer, J., & Nair, K. S. (2001). Sarcopenia. Journal of Laboratory and Clinical Medicine, 137(4), 231–243.

52. Nasreddine, Z. S., Phillips, N. A., Bédirian, V., Charbonneau, S., Whitehead, V., Collin, I.,…Chertkow, H. (2005). The Montreal Cognitive Assessment, MoCA: a brief screening tool for mild cognitive impairment. Journal of the American Geriatrics Society, 53(4), 695–699.

53. Oldfield, R. C. (1971). The assessment and analysis of handedness: the Edinburgh inventory. Neuropsychologia, 9(1), 97–113.

54. Ostry, D. J., Darainy, M., Mattar, A. A. G., Wong, J., & Gribble, P. L. (2010). Somatosensory Plasticity and Motor Learning. Journal of Neuroscience, 30(15), 5384–5393.

55. Reuter, E.-M., Pearcey, G. E. P., & Carroll, T. J. (2018). Greater neural responses to trajectory errors are associated with superior force field adaptation in older adults. Experimental Gerontology, 110, 105–117.

56. Salat, D. H. (2004). Thinning of the Cerebral Cortex in Aging. Cerebral Cortex, 14(7), 721–730.

57. Sarlegna, F. R., Malfait, N., Bringoux, L., Bourdin, C., & Vercher, J.-L. (2010). Force-field adaptation without proprioception: Can vision be used to model limb dynamics? Neuropsychologia, 48(1), 60–67.

58. Schaap, T. S., Gonzales, T. I., Janssen, T. W. J., & Brown, S. H. (2015). Proprioceptively guided reaching movements in 3D space: effects of age, task complexity and handedness. Experimental Brain Research, 233(2), 631–639.

59. Seidler, R. D., Alberts, J. L., & Stelmach, G. E. (2002). Changes in Multi-Joint Performance With Age. Motor Control, (6), 19–21.

60. Seidler-Dobrin, R. D., & Stelmach, G. E. (1998). Persistence in visual feedback control by the elderly. Experimental Brain Research, 119(4), 467–474.

61. Shadmehr, R., & Mussa-Ivaldi, F. A. (1994). Adaptive representation of dynamics during learning of a motor task. Journal of Neuroscience, 14(5), 3208–3224.

62. Shadmehr, R., Smith, M. A., & Krakauer, J. W. (2010). Error Correction, Sensory Prediction, and Adaptation in Motor Control. Annual Review of Neuroscience, 33(1), 89–108.

63. Slack, J. R., Hopkins, W. G., & Williams, M. N. (1979). Nerve Sheaths and Motoneurone Collateral Sprouting. Nature, 282, 506–507.

64. Smith, J. L., Crawford, M., Proske, U., Taylor, J. L., & Gandevia, S. C. (2009). Signals of motor command bias joint position sense in the presence of feedback from proprioceptors. Journal of Applied Physiology, 106(3), 950–958.

65. Sorock, G. S., & Labiner, D. M. (1992). Peripheral neuromuscular dysfunction and falls in an elderly cohort. American Journal of Epidemiology, 136(5), 584–591.

66. Taylor, M. M., & Creelman, C. D. (1967). PEST: Efficient Estimates on Probability Functions. The Journal of the Acoustical Society of America, 41(4), 782–787.

67. Trewartha, K. M., Garcia, A., Wolpert, D. M., & Flanagan, J. R. (2014). Fast But Fleeting: Adaptive Motor Learning Processes Associated with Aging and Cognitive Decline. Journal of Neuroscience, 34(40), 13411–13421.

68. Valdez, G., Tapia, J. C., Kang, H., Clemenson, G. D., Gage, F. H., Lichtman, J. W., & Sanes, J. R. (2010). Attenuation of age-related changes in mouse neuromuscular synapses by caloric restriction and exercise. Proceedings of the National Academy of Sciences, 107(33), 14863–14868.

69. van Beers, R. J., Sittig, A. C., & van der Gon, J. J. D. (1998). The precision of proprioceptive position sense. Experimental Brain Research, 122(4), 367–377.

70. Vindras, P., Desmurget, M., Prablanc, C., & Viviani, P. (1998). Pointing Errors Reflect Biases in the Perception of the Initial Hand Position. J Neurophysiol, 79(6), 3290–3294.

71. Wilson, E. T., Wong, J., & Gribble, P. L. (2010). Mapping proprioception across a 2D horizontal workspace. PloS One, 5(7), e11851.

72. Wingert, J. R., Welder, C., & Foo, P. (2014). Age-Related Hip Proprioception Declines: Effects on Postural Sway and Dynamic Balance. Archives of Physical Medicine and Rehabilitation, 95(2), 253–261.

73. Wolpert, D. M., Ghahramani, Z., & Jordan, M. I. (1995). An internal model for sensorimotor integration. Science, 269(5232), 1880–1882.

74. Wong, J. D., Wilson, E. T., Kistemaker, D. A., & Gribble, P. L. (2014). Bimanual proprioception: are two hands better than one? Journal of Neurophysiology, 111(6), 1362–1368.

75. Wright, M. L., Adamo, D. E., & Brown, S. H. (2011). Age-related declines in the detection of passive wrist movement. Neuroscience Letters, 500(2), 108–112.

76. Yan, J. H., Thomas, J. R., Stelmach, G. E., & Thomas, K. T. (2000). Developmental Features of Rapid Aiming Arm Movements Across the Lifespan. Journal of Motor Behavior, 32(2), 121–140.

77. Yousif, N., Cole, J., Rothwell, J., & Diedrichsen, J. (2015). Proprioception in motor learning: lessons from a deafferented subject. Experimental Brain Research, 233(8), 2449–2459.

